# Identification of a putative nuclear localization signal in maspin protein shed light into its nuclear import regulation

**DOI:** 10.1101/474379

**Authors:** Jeffrey Reina, Lixin Zhou, Marcos R.M. Fontes, Nelly Panté, Nathalie Cella

**Affiliations:** Department of Cell and Developmental Biology, Institute of Biomedical Science of University of São Paulo, São Paulo – SP, Brazil; Department of Zoology, University of British Columbia, Vancouver – BC, Canada; Department of Physics and Biophysics, Institute of Biosciences, São Paulo State University (UNESP), Botucatu - SP, Brazil

## Abstract

Maspin (SERPINB5) is a potential tumor suppressor gene with pleiotropic biological activities, including regulation of cell proliferation, death, adhesion, migration and gene expression. Several studies suggest that subcellular localization plays an essential role on maspin tumor suppression activity. In this study we investigated the molecular mechanisms underlying maspin nucleocytoplasmic shuttling. An *in vitro* nuclear-import assay using digitonin-permeabilized HeLa cells demonstrated that maspin enters the nucleus by an energy-and carrier-independent mechanism. However, previous studies indicated that maspin subcellular localization is regulated in the cell. Using a nuclear localization signal (NLS) prediction software, we identified a putative NLS in the maspin amino acid sequence. To distinguish between passive and regulated nuclear translocation, maspinNLS or the full-length protein (MaspinFL) were fused to 5GFP, rendering the construct too large to enter the nucleus passively. Unexpectedly, 5GFP-maspinNLS, but not maspinFL-5GFP, entered the nucleus of HeLa cells. Dominant-negative Ran-GTPase mutants RanQ69L or RanT24N, suppressed 5GFP-maspinNLS nuclear localization. In summary, we provide evidence that maspin translocates to the nucleus passively. In addition, we identified a peptide in the maspin protein sequence, which is able to drive a 5GFP construct to the nucleus in an energy-dependent manner.

## Introduction

Maspin (mammary serine protease inhibitor) also known as SerpinB5, is a potential tumor suppressor gene first identified in breast tissue [1]. It is now well established that maspin is expressed by most epithelia [2] and has diverse biological activities, including inhibition of tumor growth and invasion, regulation of cell adhesion, migration, apoptosis, gene transcription and oxidative stress response [3]. Maspin tumor suppressor activity is complex and appears to be cell type and tissue-context dependent. Downregulation of maspin expression has been observed in some tumor types [1,4,5], whereas others found an opposite trend [6–10]. Interestingly, increasing evidences indicate that maspin nuclear localization, rather than its level of expression, correlates with good prognostic and tumor suppression [10–12]. These data underscore the importance of understanding the molecular mechanism underlying maspin nucleocytoplasmic traffic, which can lead to new cancer therapeutic targets. Regulated nuclear transport is a signal-mediated process dependent on nuclear transporters of the Karyopherin-β superfamily (Kabβ), also called importins and exportins [13]. The 20 different human Kabβ subfamilies recognize a NLS within the cargo polypeptide. The cargo-importin complex binds to nucleoporins of the nuclear pore complex (NPC) and to the small GTPase Ran, which provides both energy and directionality to the transport. As a 42 kDa protein, maspin can potentially enter the nucleus by passive diffusion [14]. However, when maspin cDNA was transfected into mouse mammary tumor TM40D cells, maspin was found exclusively in the cytoplasm [15]. In addition, we found that a fraction of cellular maspin accumulates in the nucleus in EGF-treated MCF-10A cells and it is predominantly cytoplasmic in the mouse mammary gland [16]. Based on the observations described above, we hypothesized both passive and active/regulated mechanisms regulate maspin nucleocytoplasmic shuttling.

To test this hypothesis, we reconstituted maspin nuclear import *in vitro* using digitonin-permeabilized HeLa cells. We observed that maspin promptly translocates to the nucleus in the absence of exogenous cytosol or energy-regenerating solution, indicating that maspin enters the nucleus passively. Using the nuclear localization signal (NLS) predictor cNLS Mapper [17], we identified a putative bipartite NLS of 28 amino acids in the maspin protein sequence. In order to investigate if this sequence plays a role on active/regulated maspin nuclear import, full length and the maspin putative NLS sequence were cloned into a plasmid encoding five GFP molecules in tandem, generating maspinFL-5GFP and 5GFP-maspinNLS constructs respectively. When the corresponding proteins are expressed, it is expected that they do not passively diffuse because 5GFP is too large to passively translocate to the nucleus [14]. Surprisingly, maspin NLS, but not maspin full length, was able to drive nuclear import of the 5GFP construct, indicating that this peptide sequence can mediate an active transport to the nucleus. As active nuclear transport requires energy provided by Ran-GTPase-mediated GTP hydrolysis, we further investigate 5GFP-maspinNLS nuclear transport in the presence of the RanQ69L and RanT24N mutants, which are deficient in GTP hydrolysis or does not bind to GTP, respectively, and therefore act as dominant negative inhibitors of signal-and energy-dependent nuclear transport [18,19]. We observed that 5GFP-maspinNLS nuclear import was completely inhibited when Ran-GTPase mutant plasmids were co-transfected in HeLa cells. Herein, we propose a model in which different pools of maspin translocates by passive and active mechanisms. Part of the identified NLS is buried into maspin tridimensional structure, which may explain why it failed to drive nuclear translocation of the full length protein.

## Results

### Maspin diffuses into the nucleus of digitonin-permeabilized cells

Maspin is found in the nucleus, cytoplasm or in both compartments depending on the physiological conditions and cell-type [20]. As a 42 kDa protein, maspin may be able to translocate passively to the nucleus. In order to test this, we took advantage of the widely used nuclear import assay, where the plasma membrane is selectively permeabilized by digitonin, releasing cytoplasmic components and leaving the nuclear envelope intact [21]. HeLa cells were permeabilized with digitonin and the integrity of the nuclear envelope was tested with Texas Red-labeled 70 kDa dextran. In this assay fluorescence was detected in the cytoplasm only (Figure 1A), confirming the integrity of the nuclear envelope. Recombinant maspin or BSA covalently linked to the NLS of SV40 T antigen were conjugated to Cy3 and assayed in digitonin-permeabilized HeLa cells in the presence or absence of energy regenerating system (+ energy+ or - energy) and rabbit reticulocyte lysate (RRL), a source of cytoplasmic factors. As expected, BSA-NLS-Cy3 was promptly detected in the nuclei of digitonin-permeabilized cells in the presence of energy and RRL (Figure 1B, right hand panels), but not in the absence of them (Figure 1B, left hand panels). Interestingly, Maspin-Cy3 entered the nucleus irrespective of the presence of energy and RRL (Figure 1C). This result indicates that maspin can potentially enter the nucleus passively.

**Fig 1.**
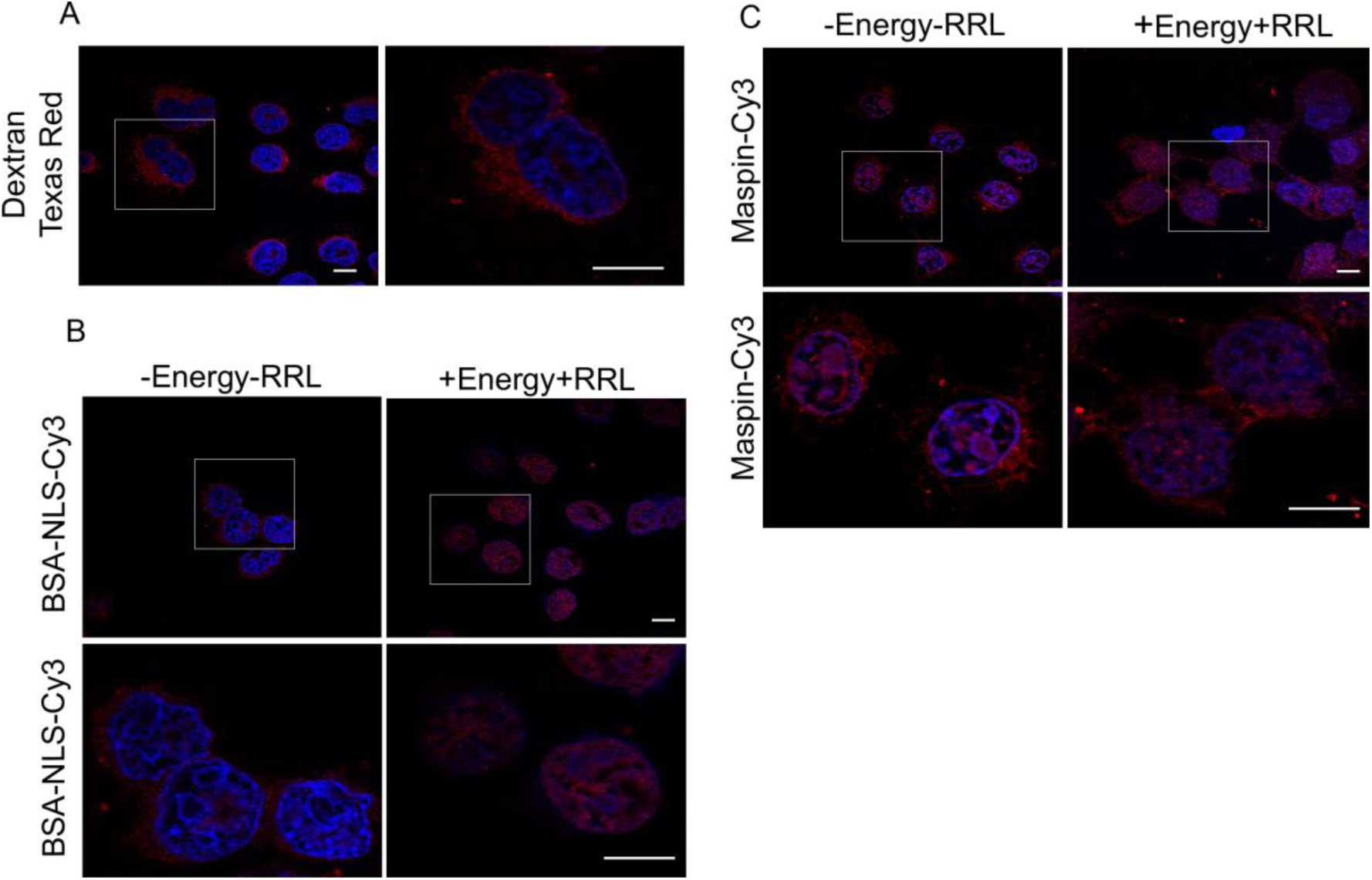
Maspin diffuses into the nucleus of digitonin-permeabilized HeLa cells. (A) HeLa cells were incubated with 70 kDa Dextran Texas Red to confirm that the nuclear membrane was intact after digitonin permeabilization; (B-C) Digitonin-permeabilized HeLa cells were incubated with BSA-NLS-Cy3 (B) or Maspin-Cy3 (C) in the presence or absence of cytosol extract (+RRL and –RRL, respectively) and with or without an energy-regenerating system (+Energy, - Energy, respectively). DAPI was used to stain the nucleus. Scale bars: 10 μm.

### Maspin amino acids 87 to 114 drives nuclear translocation of the chimeric protein 5GFP-maspinNLS

The observation that maspin can diffuse to the nucleus does not exclude the possibility that a regulated mechanism takes place. We have previously reported the existence of different maspin isoforms in the cell [16,22]. Maspin is found in different cellular compartments other than nucleus, including mitochondria [23], endoplasmic reticulum-associated vesicles [2], plasma membrane [24] and exosomes [25]. Therefore, maspin isoforms may be located in different compartments, implying a diverse regulation of subcellular localization. In addition, maspin nucleocytoplasmic distribution is determinant for its biological function as tumor suppressor and correlates with tumor prognosis. These observations underscore the importance and complexity of maspin intracellular traffic control. To determine whether maspin enters the nucleus by a carrier-dependent mechanism that requires the presence of at least an NLS on maspin, we took advantage of cNLS mapper algorithm [17]. The predicted maspin NLS sequence is the 28 amino acid sequence KLIKRLYVDKSLNLSTEFISSTKRPYAK, which was called maspinNLS. In order to distinguish between passive and regulated nuclear translocation, maspinNLS or full-length maspin (MaspinFL) were fused to five green fluorescent protein molecules in tandem (5GFP), generating proteins that are above the diffusion limit of the nuclear pore complex [14]. 5GFP-maspinNLS and maspinFL-5GFP constructs were transfected in HeLa cells. As a control, HeLa cells were also transfected with the 5GFP plasmid. The subcellular localization of the resulting chimeric proteins was assessed 24 hours post transfection using confocal laser scanning microscopy. As expected, without NLS, 5GFP was localized in the cytoplasm of the transfected cells (Figure 2A, middle panels). However, maspinNLS was able to drive nuclear transport of the 5GFP chimera protein (Figure 2A, upper panels). This suggests that maspinNLS could potentially be responsible for maspin nuclear translocation in the native molecule (Fig 2A, upper panels). Unexpectedly, MaspinFL-5GFP was not detected in the nucleus (Figure 2A, lower panels). Quantification of the nuclear to cytoplasmic fluorescence ratio (Fn/c) in these cells showed that cellular localization of maspinFL-5GFP was indistinguishable from 5GFP-transfected cells (Figure 2B). The same result was observed in transfected MCF10A cells (data not shown), a non-transformed mammary epithelial cell line, indicating that the exclusion of MaspinFL-5GFP from the nucleus is not restricted to the HeLa cell line.

**Fig 2.**
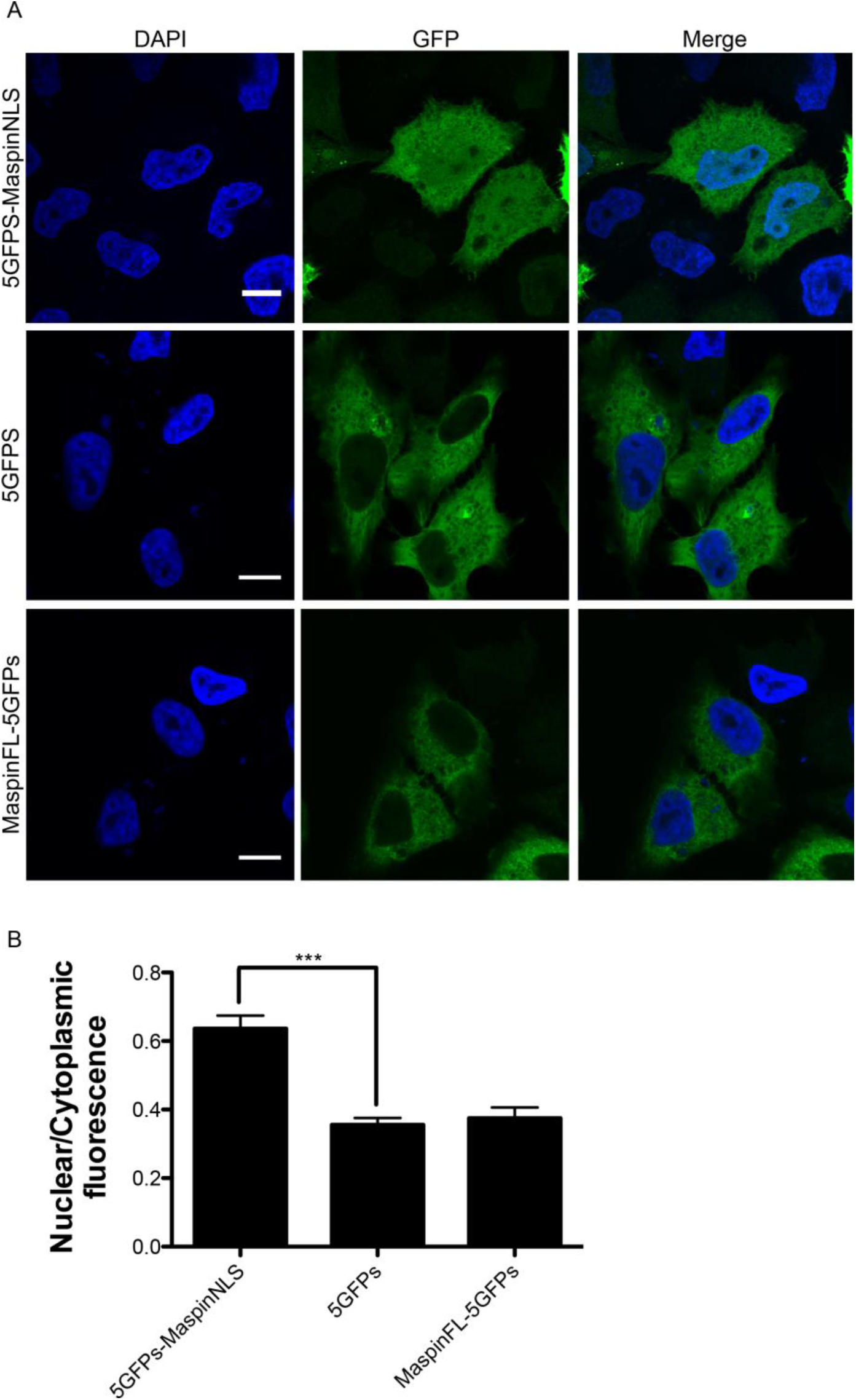
Maspin putative NLS, but not MaspinFL, fused to 5GFP induces nuclear transport of the chimera protein. (A) HeLa cells were transfected with 5GFP-MaspinNLS, MaspinFL-5GFP and 5GFP plasmids. After 24h cells were fixed and visualized with a confocal microscope. DAPI was used to stain the nucleus. Scale bars: 10 μm. (B) Quantification of nuclear to cytoplasmic fluorescence ratio from confocal microscope images. Fluorescence of 52 cells was measured for each condition. Graph bars show mean ± S.E.M. of the three different plasmids.

### Nuclear import of 5GFP-maspinNLS depends on Ran-GTPase

Active nuclear transport depends on the Ran-GTPase, although non-conventional mechanisms, which are both importin and Ran-GTP-independent do occur [26]. To further characterize the mechanism of maspin nuclear translocation, HeLa cells were cotransfected with 5GFP-maspinNLS together with plasmids encoding fluorescent wild-type (WT) Ran-GTPase, RanQ69L or RanT24N mutants. As control, cells were cotransfected with these Ran plasmids and the 5GFP plasmid, instead of the 5GFP-maspinNLS plasmid. RanQ69L cannot undergo hydrolysis and therefore it is locked in the GTP bound form, whereas RanT24N has low affinity for GTP and therefore stays always in the GDP bound form. Both mutants have been reported to inhibit Ran dependent nuclear import [27]. 5GFP-maspinNLS was detected in the nucleus when it was cotransfected with wild-type Ran-GTPase (Figure 3A, upper panels), but not when it was cotransfected with RanT24N (Figure 3A, middle panels) or RanQ69L (Figure 3A, lower panels). As expected, 5GFP subcellular localization was not affected when it was cotransfected with wild-type or Ran mutants (Figure 3B). Quantification of the nuclear to cytoplasmic fluorescence ratio (Fn/c) in these cells showed that Ran mutants significantly inhibited 5GFP-MaspinNLS translocation to the nucleus (Figure 3C). This result indicates that maspinNLS depends on a functional Ran-GTPase in order to drive nuclear translocation of the 5GFP construct.

**Fig 3.**
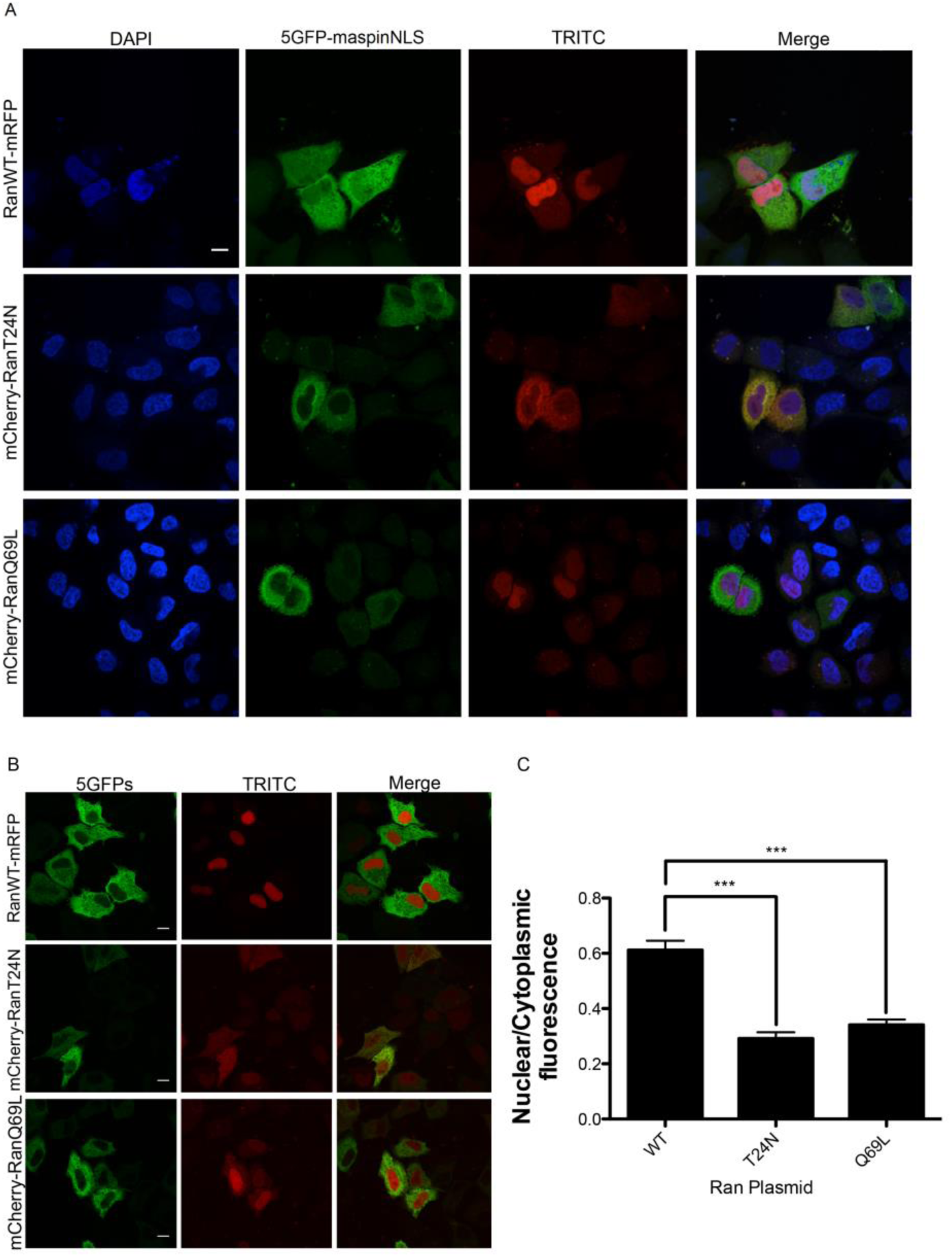
Ran mutants inhibited 5GFP-MaspinNLS nuclear import in HeLa cells. (A-B) HeLa cells were co-transfected with plasmids 5GFP-MaspinNLS (A) or 5GFP (B) plasmids and RanWT-mRFP, mcherry-RanT24N, or mCherry-RanQ69L plasmids. After 24h cells were fixed and visualized with a confocal microscope. DAPI was used to stain the nucleus. Scale bars: 10 μm. (C) Quantification of nuclear to cytoplasmic fluorescence ratio from confocal microscope images. Fluorescence of 50 cells was measured for each condition. Graph bars show mean ± S.E.M. of the three different Ran plasmids

## Discussion

Our results indicate that maspin NLS is able to transport the chimera 5GFP-maspinNLS into the nucleus. However, maspinFL-5GFP does not enter the nucleus. At this point two essential questions need to be addressed – why does not maspinNLS promote nuclear translocation of maspinFL? Which importins are involved in maspinNLS-mediated nuclear translocation? As cNLS Mapper was primarily designed to identify importin-α-dependent classical NLS [17], we first asked if the predicted NLS would interact with importin α1 (KPNA2), which is considered a general importer of cargoes bearing classical NLS [28]. Co-immunoprecipitation assays and isothermal titration calorimetry, however, did not confirm this hypothesis (data not shown). A recent method called SILAC-Tp allowed the identification of cargoes for different importins [29]. In one of this studies, maspin was ranked high among importin-11 cargoes and possibly as a transportin-1 (Kapβ2) cargo as well [30]. Interestingly, maspinNLS peptide is suitable for a PY-NLS signal (KLIKRLYVDKSLNLSTEFISST<u>**K**</u>R<u>**PY**</u>A), which is recognized by transportin-1 [31]. We are currently investigating this possibility.

We currently do not understand why maspinNLS was not able to drive maspinFL nuclear translocation, but we speculate it might be related to the localization of this peptide in maspin tridimensional structure, as part of the maspinNLS (the β-strand 2A of the β-sheet A) is buried inside the molecule (Figure 4). Furthermore, maspin is subjected to several posttranslational modification, including phosphorylation [16,32], nitrosylation [33] and acetylation [34]. We have previously observed that EGF-induced maspin phosphorylation is followed by its nuclear translocation [16]. As phosphorylation is recognized as an important regulator of nuclear translocation [35], our results support a model in which maspin NLS is somehow hidden from the nuclear transport machinery (mainly importins). Upon different stimuli (for example, member of the EGF growth factor family), maspin phosphorylation leads to a conformational modification, which may allow NLS exposure. In addition, we observed that maspin can potentially translocate passively to the nucleus, suggesting the presence of different pools of maspin in the cell, which are differentially regulated. In conclusion, our data shed light into the mechanism of maspin nuclear translocation, which may lead to a better understanding of maspin biological and tumor suppression function.

**Fig 4.**
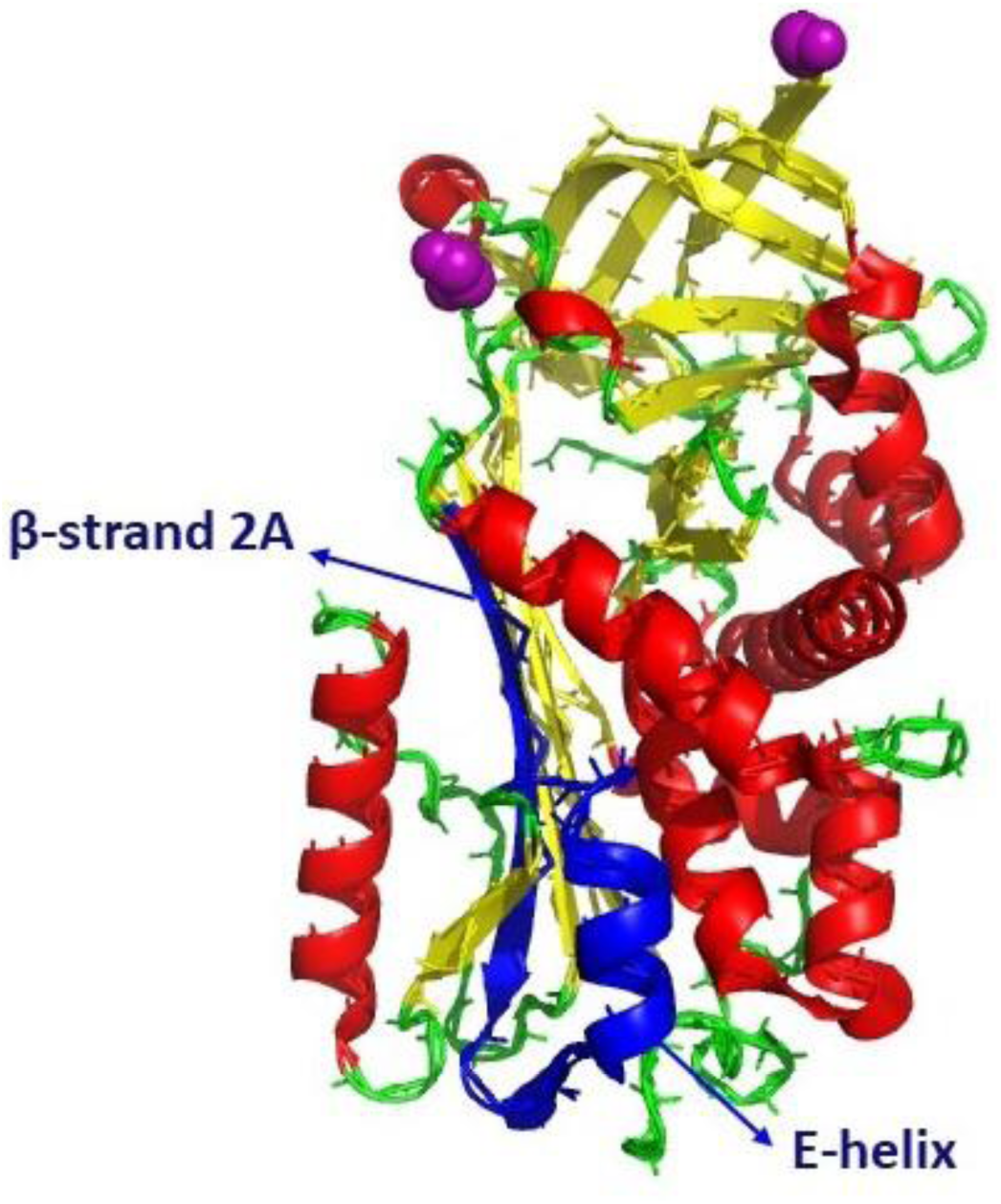
Maspin X-ray crystal structure [40]. Ribbon diagram α-helices are in red, β-strands in yellow, the RCL (reactive center loop) in purple and coils and turns are in green. The predicted NLS in maspin is shown in blue. Figure was generated with PyMOL [41].

## Materials and Methods

### Cell culture

HeLa cells were obtained from the American Type Culture Collection and were cultured in Dulbecco’s modified Eagle’s medium (DMEM) (Sigma-Aldrich) supplemented with 5% fetal bovine serum (Sigma-Aldrich), 1% penicillin-streptomycin (Cellgro), 1% L-glutamine (Cellgro) and 1% sodium pyruvate (Life Technologies). Cells were maintained at 37ºC with 5% CO_2_.

### Nuclear Import assay in Digitonin-permeabilized cells

BSA covalently attached to the NLS of SV40T antigen (CGGGPKKKRKVED) at a ratio of 5:1 (NLS:BSA) was custom made (Sigma-Genosys). Cy3 protein labeling was done with the Cy3 bis-Reactive dye pack (GE Healthcare Amersham) following manufacturer’s instructions.

HeLa cells were grown on glass coverslips until they were 40-60% confluent, washed once with phosphate buffered saline (PBS) and once with import buffer (20 mM HEPES pH 7.4, 110 mM potassium acetate, 1 mM EGTA, 5 mM sodium acetate, 2mM magnesium acetate, 2 mM DTT and 10 μg/mL protease inhibitors). For permeabilization, cells were incubated with digitonin (20 μg/ml) for 3 min at room temperature and washed three times with import buffer. Permeabilized cells were incubated with or without an energy regenerating system (0.4 mM ATP, 0.45 mM GTP, 4.5 mM phosphocreatine, 18 U/ml phosphocreatine kinase, 1.6 mg/ml BSA) and 20% cytosol extract obtained from nuclease-treated Rabbit Reticulocyte Lysates (RRL) (Promega) in the presence of 0.2 μg of 70 kDa fluorescent Dextran Texas Red (Invitrogen), 2 μg of Cy3-labeled BSA fused to SV40 NLS sequence (Sigma-Genosys), or Cy3-labeled human recombinant maspin (Peprotech) for 30 min at 37ºC. Next, the cells were washed three times with import buffer and fixed with 3% paraformaldehyde (PFA) for 10 min. Finally, the cells were washed 3 times for 5 min with PBS and mounted onto microscope slides using ProLong Diamond Antifade Mountant with DAPI (Molecular Probes). Samples were visualized using a Fluoview FV1000 confocal laser scanning microscope (Olympus).

### Maspin NLS prediction

cNLS mapper [17] was used to predict a putative NLS using human maspin amino acid sequence (UniProt identifier: P36952-1) and cut-off score of 6.0.

### Plasmids

To generate the 5GFP-MaspinNLS construct, two synthetic primers (5’-GATCCAAACTAATCAAGCGGCTCTACGTAGACAAATCTCTGAATCTTTCTACAGAGTTCATCAGCTCTACGAAGAGACCCTATGCAG-3’ and 5’ GATCCTGCATAGGGTCTCTTCGTAGAGCTGATGAACTCTGTAGAAAGATTCAGAGATTTGTCTACGTAGAGCCGCTTGATTAGTTTG-3’) were designed to include a putative bipartite NLS region in human maspin. The synthetic DNA for MaspinNLS containing adapters of the Bam HI restriction enzyme at each end were annealed, and the annealed DNA fragments were ligated to the Bam HI site at the C-terminal coding sequence of 5GFP. The construct was confirmed by DNA sequencing.

To generate MaspinFL-5GFP plasmid, human maspin cDNA was first subcloned from SERPINB5 plasmid (Origene, Catalogue number: RC224287) to pEGFP-C3 vector. The resulting maspin-GFP plasmid was used in a PCR reaction using Mas-NheI-F2 (forward) and Mas-Nhe-R-1 (reverse) primers: 5’-GAA CCG TCA GAT CCG CTA GCG CT-3’and 5’-CAC TAG CTA GCG GAG AAC AGA ATT TGC CAA AGA

AAA T-3’. The Nhe I restricted maspin PCR product was inserted into the Nhe I-digested pEGFP-GFP5 vector and ligated with T4 DNA ligase (New England Biolabs). The ligation products were transformed by heat shock into Subcloning Efficiency™ DH5α™ Competent Cells (Invitrogen) and colonies were screened for the presence of the insert. Those positive for the insert were confirmed by sequencing.

The pcDNA-RanWT-mRFP1-polyA (Addgene plasmid#59750), pTK21 (RanT24N) (Addgene plasmid# 37396) and pmCherry-C1-RanQ69L (Addgene plasmid# 30309) were gifts from Yi Zhang [36], Ian Cheeseman [37] and Jay Brenman [38], respectively.

### Transfection and Co-transfection of plasmids

HeLa cells were grown on glass coverslips in 24-well dish without antibiotics. 250 ng of DNA of each plasmid were transfected with Lipofectamine 2000 (Invitrogen). 24 h post-transfection cells were fixed with 3% PFA in PBS pH 7.4 for 10 min, washed three times with PBS and mounted onto microscope slides using ProLong Diamond Antifade Mountant with DAPI (Invitrogen). Cells were visualized with a Fluoview FV1000 confocal laser scanning microscope (Olympus).

### Quantification of nuclear import

Quantification was as described in [39]. Briefly, the fluorescence intensity of defined areas (20 x 20 pixels) was measured in the nucleus (Fn) and in the cytoplasm (Fc) using ImageJ (National Institute of Health). The mean background intensity (MB) was also measured. The ratio of nucleus to cytoplasm fluorescence intensity was calculated using the equation: Fn/c = (Fn-MB)/(Fc-MB). Data were obtained from at least 50 cells per experiment. Normal distribution was tested for each set of data using Shapiro-Wilk normality test. In the case of normal distribution, a One-way ANOVA was used to compare means between groups. In the case of non normal distribution, a Kruskal-Wallis test was used instead. To compare all groups a post-hoc Dunns test was used. All the statistical analyses were performed using GraphPad Prism and a p-value<0.05 was considered significant. All data are represented as mean ± standard error of the mean (S.E.M.).

## Acknowledgements

We are grateful to Dr. Gergely L. Lukacs (McGill University) for the kind gift of the pEGFP-GFP5 vector and to Dr. Shaker Chuck Farah (Chemistry Institute, University of São Paulo) for his protein structure explanations. This work was supported by grants from the Natural Sciences and Engineering Research Council of Canada (NSERC; RGPIN 227926-11 and RGPIN-2017-04600 to N.P.) and from São Paulo Research Foundation (FAPESP # 2013/24705-3 to N.C. and 2015/09309-0 to M.R.M.F.). Fellowships to J.R were from Coordination for the Improvement of Higher Education Personnel (CAPES - Scholarships for Foreign Students-PEC-PG) and from Emerging Leaders Of America Program (ELAP, Global Affairs Canada).

## References

1. Zou Z, Anisowicz A, Hendrix MJ, Thor A, Neveu M, et al. (1994) Maspin, a serpin with tumor-suppressing activity in human mammary epithelial cells. Science 263: 526–529.

2. Pemberton PA, Tipton AR, Pavloff N, Smith J, Erickson JR, et al. (1997) Maspin is an intracellular serpin that partitions into secretory vesicles and is present at the cell surface. J Histochem Cytochem 45: 1697–1706.

3. Bailey CM, Khalkhali-Ellis Z, Seftor EA, Hendrix MJ (2006) Biological functions of maspin. J Cell Physiol 209: 617–624.

4. Reddy KB, McGowen R, Schuger L, Visscher D, Sheng S (2001) Maspin expression inversely correlates with breast tumor progression in MMTV/TGF-alpha transgenic mouse model. Oncogene 20: 6538–6543.

5. Heighway J, Knapp T, Boyce L, Brennand S, Field JK, et al. (2002) Expression profiling of primary non-small cell lung cancer for target identification. Oncogene 21: 7749–7763.

6. Maass N, Hojo T, Ueding M, Luttges J, Kloppel G, et al. (2001) Expression of the tumor suppressor gene Maspin in human pancreatic cancers. Clin Cancer Res 7: 812–817.

7. Smith SL, Watson SG, Ratschiller D, Gugger M, Betticher DC, et al. (2003) Maspin - the most commonly-expressed gene of the 18q21.3 serpin cluster in lung cancer-is strongly expressed in preneoplastic bronchial lesions. Oncogene 22: 8677–8687.

8. Umekita Y, Yoshida H (2003) Expression of maspin is up-regulated during the progression of mammary ductal carcinoma. Histopathology 42: 541–545.

9. Bieche I, Girault I, Sabourin JC, Tozlu S, Driouch K, et al. (2003) Prognostic value of maspin mRNA expression in ER alpha-positive postmenopausal breast carcinomas. Br J Cancer 88: 863–870.

10. Mohsin SK, Zhang M, Clark GM, Craig Allred D (2003) Maspin expression in invasive breast cancer: association with other prognostic factors. J Pathol 199: 432–435.

11. Dietmaier W, Bettstetter M, Wild PJ, Woenckhaus M, Rummele P, et al. (2006) Nuclear Maspin expression is associated with response to adjuvant 5-fluorouracil based chemotherapy in patients with stage III colon cancer. Int J Cancer 118: 2247–2254.

12. Sood AK, Fletcher MS, Gruman LM, Coffin JE, Jabbari S, et al. (2002) The paradoxical expression of maspin in ovarian carcinoma. Clin Cancer Res 8: 2924–2932.

13. Cautain B, Hill R, de Pedro N, Link W (2015) Components and regulation of nuclear transport processes. FEBS J 282: 445–462.

14. Wang R, Brattain MG (2007) The maximal size of protein to diffuse through the nuclear pore is larger than 60kDa. FEBS Lett 581: 3164–3170.

15. Zhang W, Shi HY, Zhang M (2005) Maspin overexpression modulates tumor cell apoptosis through the regulation of Bcl-2 family proteins. BMC Cancer 5: 50.

16. Tamazato Longhi M, Magalhaes M, Reina J, Morais Freitas V, Cella N (2016) EGFR Signaling Regulates Maspin/SerpinB5 Phosphorylation and Nuclear Localization in Mammary Epithelial Cells. PLoS One 11: e0159856.

17. Kosugi S, Hasebe M, Tomita M, Yanagawa H (2009) Systematic identification of cell cycle-dependent yeast nucleocytoplasmic shuttling proteins by prediction of composite motifs. Proc Natl Acad Sci U S A 106: 10171–10176.

18. Dickmanns A, Bischoff FR, Marshallsay C, Luhrmann R, Ponstingl H, et al. (1996) The thermolability of nuclear protein import in tsBN2 cells is suppressed by microinjected Ran-GTP or Ran-GDP, but not by RanQ69L or RanT24N. J Cell Sci 109 (Pt 6): 1449–1457.

19. Palacios I, Weis K, Klebe C, Mattaj IW, Dingwall C (1996) RAN/TC4 mutants identify a common requirement for snRNP and protein import into the nucleus. J Cell Biol 133: 485–494.

20. Bodenstine TM, Seftor RE, Khalkhali-Ellis Z, Seftor EA, Pemberton PA, et al. (2012) Maspin: molecular mechanisms and therapeutic implications. Cancer Metastasis Rev 31: 529–551.

21. Adam SA, Marr RS, Gerace L (1990) Nuclear protein import in permeabilized mammalian cells requires soluble cytoplasmic factors. J Cell Biol 111: 807–816.

22. Tamazato Longhi M, Cella N (2012) Tyrosine phosphorylation plays a role in increasing maspin protein levels and its cytoplasmic accumulation. FEBS Open Bio 2: 93–97.

23. Latha K, Zhang W, Cella N, Shi HY, Zhang M (2005) Maspin mediates increased tumor cell apoptosis upon induction of the mitochondrial permeability transition. Mol Cell Biol 25: 1737–1748.

24. Sheng S, Carey J, Seftor EA, Dias L, Hendrix MJ, et al. (1996) Maspin acts at the cell membrane to inhibit invasion and motility of mammary and prostatic cancer cells. Proc Natl Acad Sci U S A 93: 11669–11674.

25. Dean I, Dzinic SH, Bernardo MM, Zou Y, Kimler V, et al. (2017) The secretion and biological function of tumor suppressor maspin as an exosome cargo protein. Oncotarget 8: 8043–8056.

26. Wagstaff KM, Jans DA (2009) Importins and beyond: non-conventional nuclear transport mechanisms. Traffic 10: 1188–1198.

27. Klebe C, Bischoff FR, Ponstingl H, Wittinghofer A (1995) Interaction of the nuclear GTP-binding protein Ran with its regulatory proteins RCC1 and RanGAP1. Biochemistry 34: 639–647.

28. Kohler M, Speck C, Christiansen M, Bischoff FR, Prehn S, et al. (1999) Evidence for distinct substrate specificities of importin alpha family members in nuclear protein import. Mol Cell Biol 19: 7782–7791.

29. Kimura M, Thakar K, Karaca S, Imamoto N, Kehlenbach RH (2014) Novel approaches for the identification of nuclear transport receptor substrates. Methods Cell Biol 122: 353–378.

30. Kimura M, Morinaka Y, Imai K, Kose S, Horton P, et al. (2017) Extensive cargo identification reveals distinct biological roles of the 12 importin pathways. Elife 6.

31. Lee BJ, Cansizoglu AE, Suel KE, Louis TH, Zhang Z, et al. (2006) Rules for nuclear localization sequence recognition by karyopherin beta 2. Cell 126: 543–558.

32. Narayan M, Mirza SP, Twining SS (2011) Identification of phosphorylation sites on extracellular corneal epithelial cell maspin. Proteomics 11: 1382–1390.

33. Lam YW, Yuan Y, Isaac J, Babu CV, Meller J, et al. (2010) Comprehensive identification and modified-site mapping of S-nitrosylated targets in prostate epithelial cells. PLoS One 5: e9075.

34. Weinert BT, Scholz C, Wagner SA, Iesmantavicius V, Su D, et al. (2013) Lysine succinylation is a frequently occurring modification in prokaryotes and eukaryotes and extensively overlaps with acetylation. Cell Rep 4: 842–851.

35. Nardozzi JD, Lott K, Cingolani G (2010) Phosphorylation meets nuclear import: a review. Cell Commun Signal 8: 32.

36. Inoue A, Zhang Y (2014) Nucleosome assembly is required for nuclear pore complex assembly in mouse zygotes. Nat Struct Mol Biol 21: 609–616.

37. Kiyomitsu T, Cheeseman IM (2012) Chromosome- and spindle-pole-derived signals generate an intrinsic code for spindle position and orientation. Nat Cell Biol 14: 311–317.

38. Kazgan N, Williams T, Forsberg LJ, Brenman JE (2010) Identification of a nuclear export signal in the catalytic subunit of AMP-activated protein kinase. Mol Biol Cell 21: 3433–3442.

39. Wu WW, Sun YH, Pante N (2007) Nuclear import of influenza A viral ribonucleoprotein complexes is mediated by two nuclear localization sequences on viral nucleoprotein. Virol J 4: 49.

40. Law RH, Irving JA, Buckle AM, Ruzyla K, Buzza M, et al. (2005) The high resolution crystal structure of the human tumor suppressor maspin reveals a novel conformational switch in the G-helix. J Biol Chem 280: 22356–22364.

41. Schrodinger, LLC (2015) The PyMOL Molecular Graphics System, Version 1.8.

